# Single-cell Electrochemical Aptasensor Array

**DOI:** 10.1101/2023.03.16.532912

**Authors:** Shuo Li, Yannick Coffinier, Chann Lagadec, Fabrizio Cleri, Katsuhiko Nishiguchi, Akira Fujiwara, Soo Hyeon Kim, Nicolas Clément

## Abstract

Despite several demonstrations of electrochemical devices with limits of detection (LOD) of 1 cell/mL, the implementation of single-cell bioelectrochemical sensor arrays has remained elusive due to the challenges of scaling up. In this study, we show that the recently introduced nanopillar array technology combined with redox-labelled aptamers targeting epithelial cell adhesion molecule (EpCAM) is perfectly suited for such implementation. Combining nanopillar arrays with microwells determined for single cell trapping directly on the sensor surface, single target cells are successfully detected and analyzed. This first implementation of a single-cell electrochemical aptasensor array, based on Brownian-fluctuating redox species, opens new opportunities for large-scale implementation and statistical analysis of early cancer diagnosis and cancer therapy in clinical settings.

**For Table of Contents only:** 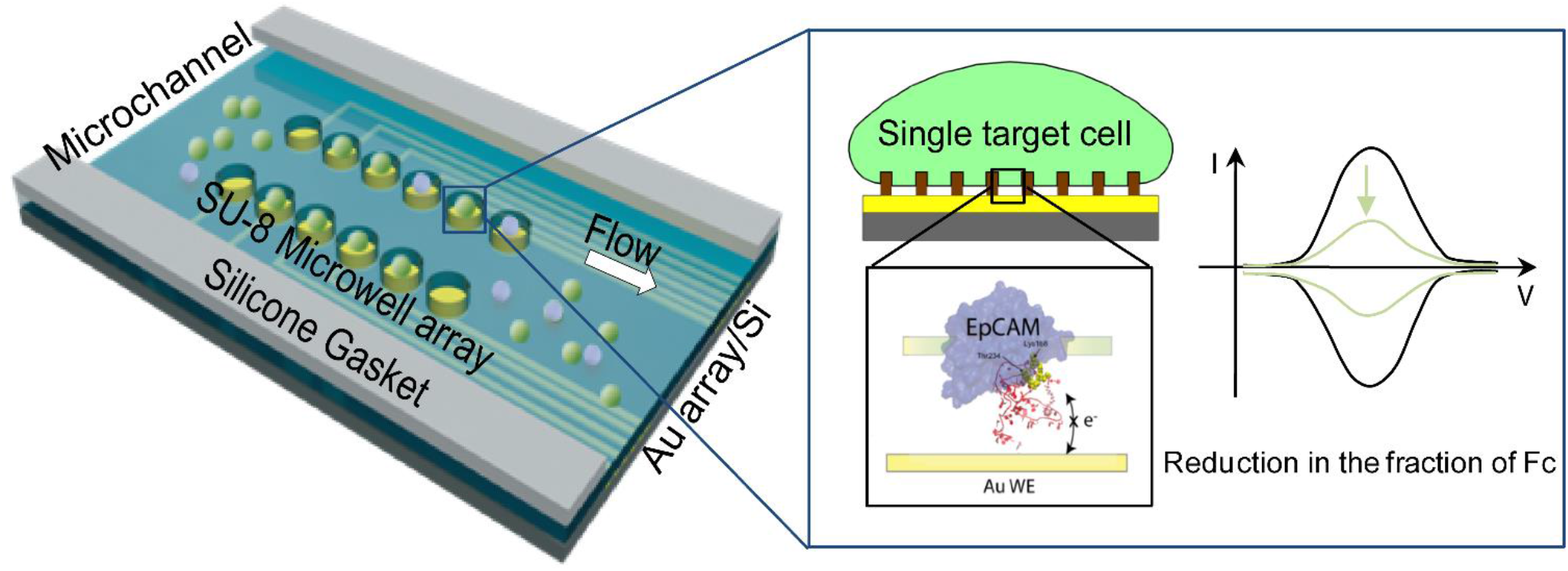

---

The need for single cell detection and analysis techniques has increased in the past decades because of the heterogeneity of individual living cells, which increases the complexity of the pathogenesis of malignant tumors.^1-3^ In the search for early cancer detection, high-precision medicine and therapy, the technologies most used today for sensitive detection of target analytes and monitoring the variation of these species are mainly including two types. One is based on the identification of molecular differences at the single-cell level, such as flow cytometry, fluorescence-activated cell sorting, next generation proteomics, lipidomic studies,^4-7^ another is based on capturing or detecting single tumor cells from fresh or fixed primary tumors and metastatic tissues, and rare circulating tumors cells (CTCs) from blood or bone marrow, for example, di-electrophoresis technique,^8^ microfluidic based microposts chip,^9^ electrochemical (EC) approach. Compared to other methods, EC sensors have the merits of easy operation, high sensitivity, and portability.^10-19^ For example, a recent technique using rolling circle amplification with aptamers could successfully demonstrate a limit of detection (LOD) as low as 5 CTC cells in whole blood (with 200 μL of the sample used for incubation).^20^ Another approach using quantum dots as sensing probe attained a LOD of 2 cells/mL in human serum.^21^ Very recently, PEDOT:PSS organic electrochemical transistors have shown the potential for electrical cell-substrate impedance sensing down to single cell.^22, 23^ However, despite various demonstrations of low LOD including aptamer sensors,^21, 24, 25^ arrayed EC sensors for detecting single-cell have not been demonstrated. Recently, we have introduced a new technique based on 20-nm-thick nanopillars array to support cells and keep them at ideal recognition distance for redox-labelled aptamers grafted on the surface. The key advantages of this technology are not only to suppress the false positive signal arising from the pressure exerted by all (including non-target) cells pushing on the aptamers by downward force, but also to stabilize the aptamer at the ideal hairpin configuration thanks to a confinement effect.^26^ With the first implementation of this technique, a LOD of 13 cells (with 5.4 μL of cell suspension) was estimated.

In this work, the nanosupported cell technology using redox-labelled aptasensors has been pushed forward and fully integrated into a single-cell electrochemical aptasensor array. To reach this goal, the LOD has been reduced by more than one order of magnitude by suppressing parasitic capacitive electrochemical signals through minimizing the sensor area and localizing the cells. A statistical analysis at the single-cell level is demonstrated for the recognition of cancer cells. The future of this technology is discussed and the potential for scaling over millions of electrodes, thus pushing further integration at sub-cellular level is highlighted. A description of the experimental methods is reported in the Supplementary Materials.

Figure 1 shows a simple description of the operation of this single-cell EC aptasensor and the configuration of the proposed microfluidic system. The device is assembled on micrometric gold working electrodes; an active biorecognition monolayer, composed of tethered ferrocene (Fc)-labelled ssDNA SYL3C aptamers^24, 25^ immobilized on the gold surfaces for the efficient and specific recognition of the epithelial cell adhesion molecules (EpCAMs), and the backfilling of oligoethylene glycol (OEG) molecules for mitigating nonspecific adsorption and stabilizing aptamers; a regular array of hydrogen-silsesquioxane (HSQ) nanopillars fabricated by high-speed e-beam lithography,^26, 28, 29^ and SU-8 microwells for gravitationally trapping single-cell (the optical image and dimensions of the SU-8 microwells are shown in Figure S1). A simple description of the fabrication process is depicted in Figure 1c (more details are depicted in Figure S2). The choice of the dimensions of HSQ nanopillars was discussed in the previous work.^26^ The atomic force microscopy (AFM) image shown in Figure 1d presents the HSQ nanopillars with a diameter of 200 nm, a height of 19.7 nm, pitch of 500 nm (a larger scale of the AFM of HSQ nanopillars that are well aligned on the Au microelectrode is shown in Figure S3); this configuration aimed to keep the cell at a distance *z*_*gap*_ of a few nanometers above the sensor surface, which is small enough to enable an interaction between the SYL3C aptamers and EpCAMs (Figure 1b), this *z*_*gap*_ has been estimated to be around 5 nm based on different nanopillars height, upon consideration of cells deformation.^26^

**Figure 1.**
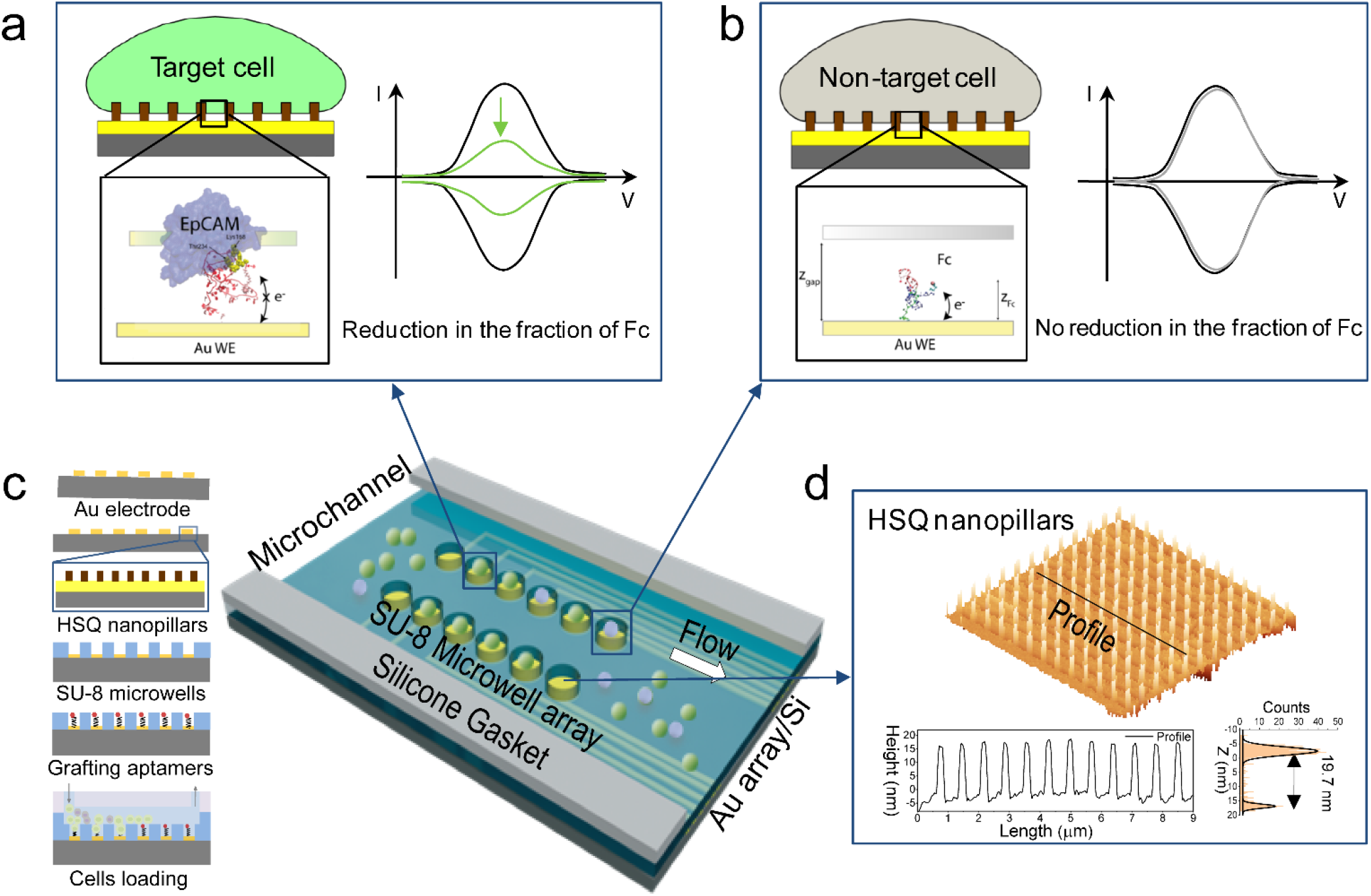
(a) Schematic representation of the nanopillars device used for the molecular recognition of EpCAM on a target cell by the SYL3C aptamer, and the relative CVs without cell (black) and with a target cell (green). (b) Schematic representation of the device composed of a gold electrode with nano-supported a non-target cell, a tethered SYL3C-Fc aptamer, as well as the relative CVs without cell (black) and with a non-target cell (grey). (c) Schematic to show the principle of electrochemical aptasensors for single cell analysis. (d) AFM image (10 μm × 10 μm) and AFM cross-section of the HSQ nanopillars.

When the cells are seeded in the microfluidic device and incubated for a short time, each single cell will be gravitationally trapped in the microwell. In the presence of a target cell, the aptamer can partly bond to the EpCAMs, and the fluctuations become more constrained. During the molecular recognition interaction, a number of H-bonds are formed between the second hairpin of the aptamer and a selected set of EpCAM residues, notably the Lys168 and the Thr234 (Figure 1b). Since the probability of charge transfer is proportional to the exponential of the distance (*z*_*gap*_), the interaction of aptamer and EpCAM drastically reduces the electron transfer rate, and the expected decrease of current is shown in Figure 1a. Therefore, the peak amplitude (*I*_*peak*_) decrease can be related statistically to the percentage of aptamers interacting with the cell, a process that is thermally activated by the formation of H-bonds. In contrast, when there is no cell or in the presence of nontarget cell, the aptamers move freely, the Fc moves in a reproducible way on long enough time constants (>>10 microseconds). In this way, electron transfer between the surface and redox markers is allowed, therefore, no reduction of the fraction of Fc is expected, *i. e*. the cyclic voltammetry curves (CVs) remain the same with and without a non-target cell (Figure 1a). The area of the voltammogram (related to either oxidation or reduction) represents the total charge transferred per cycle times the sweep rate. Therefore, it is directly related to the fraction of the aptamers on the surface that can effectively transfer the charge, and the *I*_*peak*_ is the key parameter to monitor. If the sweep rate is faster than the electron transfer or molecular diffusion rates, the shape of the voltammogram asymmetrically varies according to the oxidation and reduction peak potential.^30-32^

To evaluate the enhanced sensitivity of aptasensor with nano-supported cancer cells, two types of devices were developed by reducing the sensing area from a large scale (36mm^2^) to a smaller scale (∼700 μm^2^) specifically designed for targeting single cancer cells. Figure 2a and 2b show the fluorescent images and the associated CVs in the absence and presence of cells, respectively, utilizing the large surface sensing area. Here the lymphoma cell line Ramos was used as control experiment, which does not express Ep-CAM.^33^ The pancreatic cancer cells (Capan-2) were detected as target cells, which have a high level of EpCAM expression.^34^ The cell viability, cell number and size were determined using a Petroff-Hausser Vcell counting chamber before the detection, as shown in Figure S4. We observed a large shift in *I*_*peak*_ with 65% decrease in the presence of Capan-2 labelled with green color (Figure 2a) while no change for the *I*_*peak*_ in the control experiment, here Ramos are labelled with red color (Figure 2b), showing the effective molecular recognition by such an aptasensor as reported previously.^21^ However, in the nanopillar configuration, the LOD can be attributed to the nonfaradic current that is proportional to the sensor area and independent of the number of cells on the sensor. To reach single-cell LOD, miniaturization is helpful as the total electrode area without cells is suppressed which reduces the “parasitic” electrical double layer capacitance. Using this configuration of microelectrodes combined with microwells, single cells were selectively detected. The florescence microscopy image in Figure 2c shows single Capan-2 cell that was trapped in each microwell (a full fluorescence image of the whole one chip with 20 devices is shown in Figure S5) and the corresponding CVs, as expected, a decrease of *I*_*peak*_ (63%) was observed, the absolute *I*_*peak*_ value (figures 2c, 2d) for single-cell detection is proportionally reduced compared to the large sensing surface (figures 2a, 2b), which is in the tens of pico-Amperes. To obtain statistical results from this single-cell detection device, the CV signal was carried out only for the sensors that contained a cell in the microwell. A statistical analysis of oxidation *I*_*peak*_ of 100 devices (6 chips) for detecting a single target cell is depicted in Figure 2e. The relative standard deviation (RSD%), defined as the standard deviation divided by the means of oxidation *I*_*peak*_ for both cases (without a cell and with one target cell), are calculated as 2.8% and 10.7%, respectively. The distribution of the reduction *I*_*peak*_ and original CVs are shown in Figure S6. In addition, statistics of single non-target cells (Ramos) in 7 chips and their relative CVs are performed in Figure 2d, 2f and Figure S6. The RSD% of oxidation *I*_*peak*_ in figure 2f without a cell and with one non-target cell are estimated as 1.6% and 1.7%, respectively. Therefore, it can be concluded that a significant relative fluctuation is only observed in the presence of the target cell. It is interesting to note that the current suppression for the target cells is proportional to the cell area. The relative dispersion can be attributed to the cell diameter dispersion and microwell diameter, which is reasonable considering a microwell diameter of 30 μm, as well as the fact that the cell diameter of Capan-2 has a mean of 16.02 μm and standard deviation of 3 μm (shown in Figure S4). The results indicate that this aptasensor can be used for single-cell detection and the molecular recognition analysis at the single-cell level.

**Figure 2.**
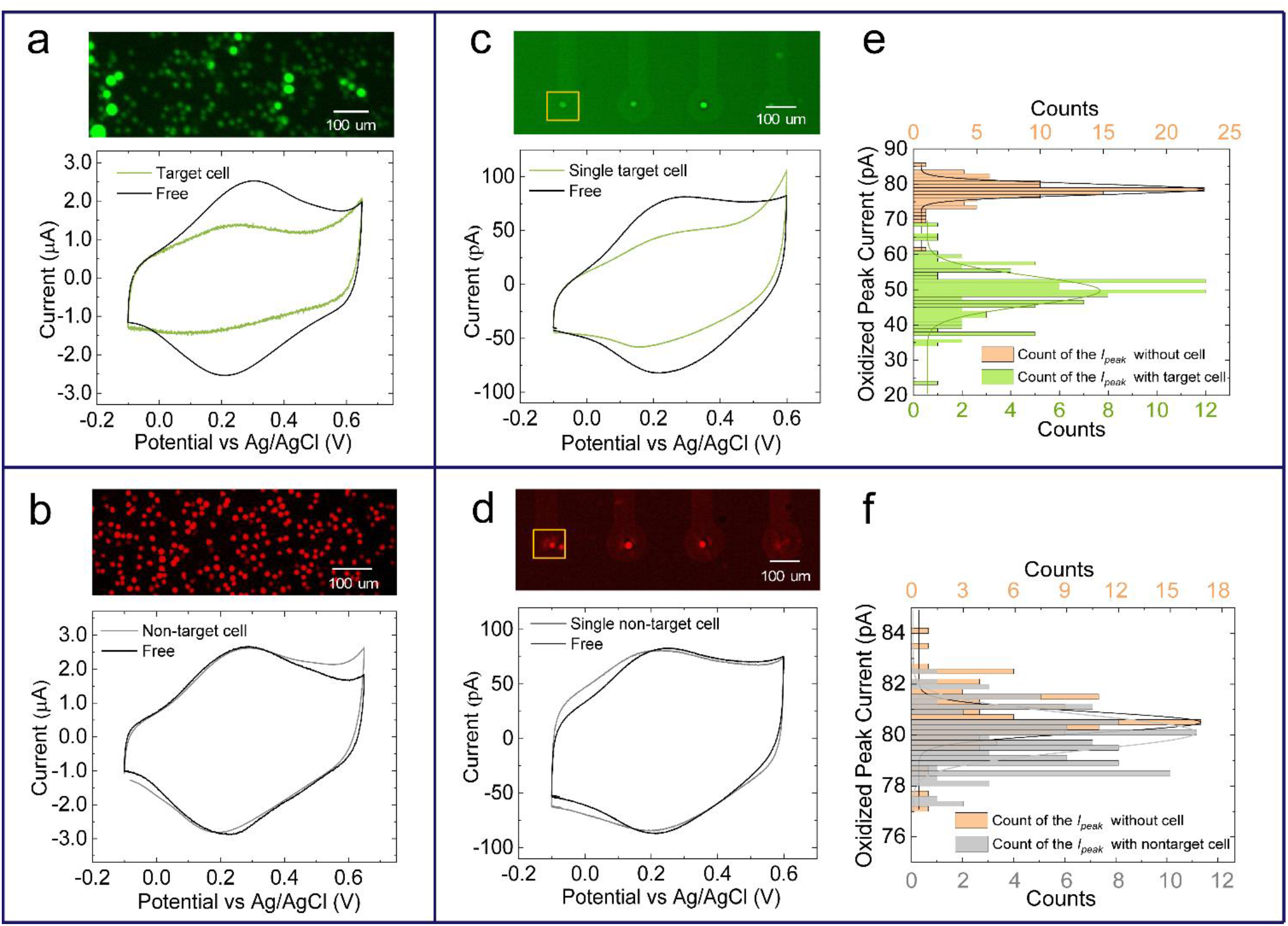
(a) Fluorescent image of target cell on the device with large-area surface and CVs (scan rate *v* = 0.5 V/s) in the presence and absence of target cells, respectively. (b) Florescent image of non-target cell on the device with large-area surface and CVs (*v* = 0.05 V/s) of a device in the presence and absence of non-target cells, respectively. (c) Fluorescent image of one target cell on the device with small-area surface and CVs (*v* = 0.5 V/s) in the presence and absence of one target cell, respectively. (d)Fluorescent image of one non-target cell on the device with small-area surface and CVs (*v* = 0.05 V/s) in the presence and absence of one non-target cell, respectively. (e) Statistics of oxidation *I*_*peak*_ of 100 devices (one cell on the surface of one device) for detecting single target cell. (f) Statistics of oxidation *I*_*peak*_ of 100 devices (one cell on the surface of one device) for detecting single non-target cell.

To further demonstrate the selectively of the aptasensor device. Figure 3 shows the results of the device working with mixed-cells (50% Capan-2 cells labelled with green color, and 50% Ramos cells labelled with red color). After loading the mixed cells, they were incubated for 30 min (Figure 3a), fresh PBS was flowed to remove the non-trapped cells. By checking the Fluorescence microscopy, it is clear to see that one target cell is on sensor 1 and one non-target cell is trapped on sensor 2, as depicted in Figure 3b. The relative CVs with different signal change for the single target cell and single non-target cell in Figures 3c, 3d, indicating the good selectivity of the device in mixed cells. This result showing the promising for granting multi-molecular studies with nanoelectrodes, for instance, interdigitated nanoelectrode,^11^ nanowires based aptasensors.^35^ Interestingly and significantly, the recent work of atto-Ampere electrochemistry for redox-labelled molecules would be directly beneficial to this EC aptasensor with nano-supported single-cell approach for subcellular sensing.^36^ Furthermore, this single-cell array shows the new opportunity for large-scale implementation and statistical analysis of clinical samples, combing our cells filtration method,^8^ and the present technology implemented in a 4 inch wafer with 4×10^5^ electrodes that could be used for detecting 1-10 CTC in (1-4)× 10^3^ cells.

**Figure 3.**
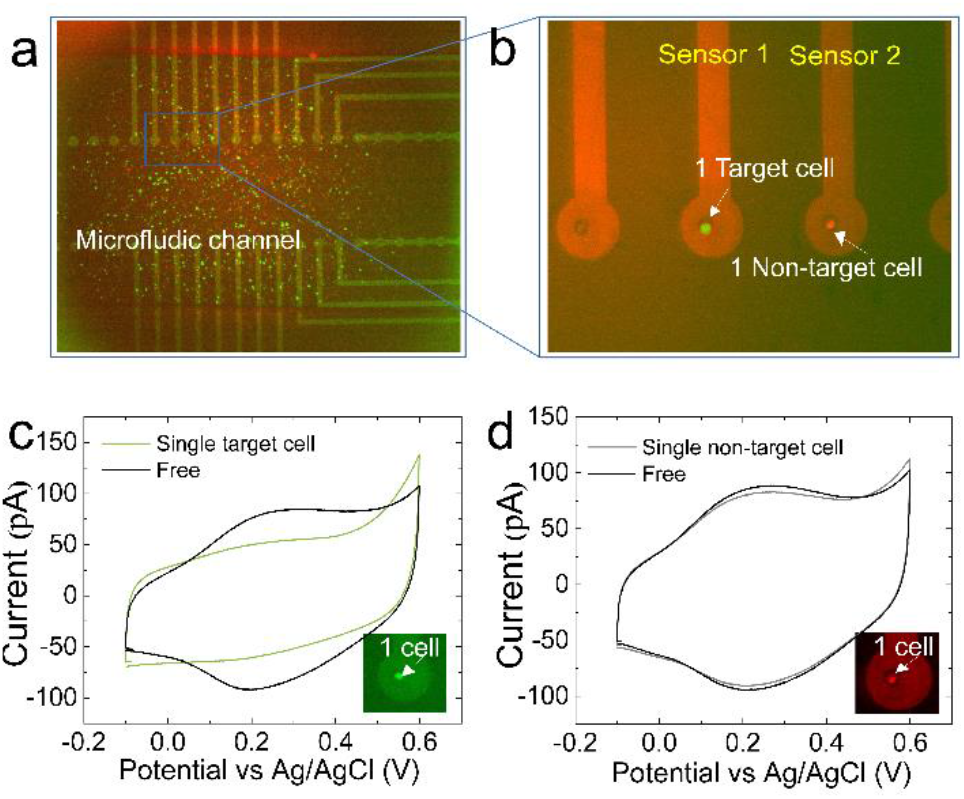
(a) Fluorescent image of mixed cells loaded on the device. (b) A zoomed-in section of (a) shows the single target cell (green) and non-target cell (red) trapped in the microelectrode. (c) CVs (*v* = 0.05 V/s) in the presence and absence of one target cell, respectively. (d) CVs (*v* = 0.05 V/s) of a device in the presence and absence of one non-target cell, respectively.

In conclusion, an EC aptasensor array with nanopillars was demonstrated for single cancer cell detection. The fabrication approach with nanogap suppresses cell downward force issue on the electrode, that allows the aptamer to move freely and thereby facilitating the process of molecular recognition. The statistical analysis for non-target single cell and target single cell shows ultra-sensitivity, selectivity, stability, and reproducibility of the device. This breakthrough method is compatible with single-cell trapping techniques such as dielectrophoresis (DEP)^8^ and shows outstanding potential for large scale implementation of single-cell arrays, it also pushes forward perspectives of using EC devices for further miniaturization at subcellular or single-molecule analysis.

## ASSOCIATED CONTENT

### Supporting Information

Experimental and methods, illustration of fabrication of the device, microscopy images of the SU-8 microwells, cell information, additional AFM, additional Fluorescence microscopy images, additional CVs.

## AUTHOR INFORMATION

### Author Contributions

The manuscript was written through contributions of all authors. All authors have given approval to the final version of the manuscript.

### Funding Sources

This work was supported by JSPS Core-to Core Program (JPJSCCA20190006), and the projects “Agence Nationale de la Recherche” (ANR) SIBI(ANR-19-CE42-0011-01), CNRS MITI BIOSTAT, and EU-Attract2 UNICORN-DX project.

### Notes

The authors declare no competing financial interest.

## ACKNOWLEDGMENT

The authors thank Dr. E. Lebrasseur, Dr. Fujiwara from Takeda CR (Utokyo) for their help and advice with device fabrication, Dr. S. Grall and Dr. L. Jalabert for discussion of the measurement.

## SI-1. Experimental and Methods Section

### 1.1 Fabrication of the single-cell detection device

#### Fabrication of microelectrode array

The working microelectrode (Φ = 100 μm) was fabricated by using laser lithography (HEIDELBERG DWL66+). First, 100 nm SiO_2_ was thermal deposited on 4-inch silicon wafer by using sputtering (Sputter Ulvac SIH-450). Then positive optical resist SPIR 3251 was spin-coated with a speed of 3000 rpm for 30 s and baked at 100 °C for 2 min. The baked substrate was then loaded into the laser drawing machine and finished writing by setting proper parameters. After writing, post exposure baking process (110 °C for 1 min 30 s) was processed and followed to immerse the substrate into NMD-3 (2.38% tetramethylammonium hydroxide) for 2 min 30 s and dried it with nitrogen gas, finally, the substrate was post baked at 120 °C for 2 min to get the desired pattern. Next, the 2 nm Ti and 50 nm Au layers were deposited on the substrate by using sputtering equipment (Sputter Shibaura CFS-4EP-LL i-Miller), the Au microelectrode pattern was obtained in the end by finishing lift-off process, that immersed the sample in acetone for 4-6 hours and subsequentially cleaned with isopropanol and deionized water for 5 min, respectively.

#### Alignment fabrication of hydrogen silsesquioxane (HSQ) nanopillars

HSQ (Dow Corning XR-1541) negative electron beam (e-beam) resist was used to fabricate the nanopillars by using e-beam lithography; Here the Advantest F7000s-VD02 e-beam equipment was used. The HSQ resin in a carrier solvent of methyl isobutyl ketone was spin-coated on the microelectrode sample with a speed of 5000 rpm for 60 s and then baked at 150 °C for 2 min on a hot plate. The cured sample was then loaded into the e-beam machine and was exposed with dose of 500 μC/cm^2^ by using the on-the-fly (OTF) mode with alignment tool operation. The exposed sample was finally immersed in the alkaline developer NMD-3 with agitation of 25 rpm for 5 min and then cured at 150 °C for 5 min to obtain the desired nanopillar array. The design of the HSQ nanopillar dimensions was in accordance with a previous report.^1^

#### Fabrication of SU-8 microwells

SU-8 microwells for gravitationally trapping single cell were aligned on the microelectrodes (with HSQ nanopillars) by using UV lithography (mask aligner PEM-800). SU8-3010 resist was spin-coated on the cleaned substrate with a speed of 1000 rpm for 30s and soft baked on the hot plate at 95°C for 15 min. The baked sample was exposed for 13s with exposure energy of 1.99×20 mJ/cm^2^ by using a photomask and aligner tool. The sample was then baked at 65°C for 1 min and 95°C for 5 min, followed by development in SU-8 developer for 5 min with agitation of 35rpm. Then sample was finally rinsed with isopropanol and deionized water, dried with nitrogen gas to get the microwells with dimension of ∼20 μm height and ∼30 μm in diameter. The sample was stored in the vacuum drying box before use.

### 1.2 Preparation of self-assembled DNA/oligoethylene glycol (OEG) mixed monolayers

The above fabricated chip was cleaned in acetone under sonication for 5 min, followed by rinsing with isopropanol and deionized water, and finally dried with nitrogen gas. Then, the cleaned chip was immersed in the freshly prepared 1 μM Fc-SYL3C (ssDNA)-SH solution. The functionalized aptamer sequence is as follows: ferrocene-5′-CAC TAC AGA GGT TGC GTC TGT CCC ACG TTG TCA TGG GGG GTT GGC CTG-(CH_2_)_3_ - 3′-SH (purchased from Biomers). The control of the density of molecule was discussed in the previous work.^2^ After that, the sample was immersed in a solution of 1 mM OEG-SH in pure ethanol (containing an ethylene glycol repeat unit HS(CH_2_)_3_(OCH_2_CH_2_)_6_OCH_2_COOH, purchased from Prochimia) for 1 h and rinsed with ethanol for 10 s to obtain the mixed self-assembled monolayer surface. The sample was cleaned with a solution of Tris EDTA (1X) contains 0.05% Tween 20 for 10 s before use.

### 1.3 Cyclic Voltammetry (CV) measurements

The VersaSTAT 4 (Princeton Applied Research) instrument was used for electrochemical measurements. A commercial flow cell (SEC-3F Spectroelectrochemical flow cell, including Ag/AgCl reference electrode and platinum counter electrode. purchased from ALS Global) was used to set the sample and connect the output of the equipment to perform the measurements. CV mode was selected for demonstrating the formation of the sensing probes and cell detection, Phosphate-buffered saline (PBS) buffer solution is used as electrolyte. During the CV experiments, the electrical potential ranged from -0.1 V to -0.6 V vs. Ag/AgCl with scan rate *v* of 0.05 V/s. To determine the density of the molecules on the electrode surface, the values were calculated by subtracting the background obtained from the capacitance.

### 1.4 Cell culture

Capan-2 cells (human pancreatic adenocarcinoma cell line) were purchased from cells lines service (CLS) and Ramos cells (Burkitt lymphoma cell line) were purchased from JCRB Cell Bank. Capan-2 and Ramos cells were cultured in 16 mm dish with RPMI 1640 medium supplemented with 10% fetal bovine serum and 100 mg/mL penicillin-streptomycin in a 5% CO_2_ atmosphere at 37 °C. The cells were collected and separated from the medium by centrifugation at 1000 rpm and 4 °C for 5 min, and then resuspended in a sterile PBS solution (pH 7.4) to obtain a homogeneous cell suspension. The cell suspension was freshly prepared just before incubation with the device. The cell size and cell density were determined using a Petroff-Hausser Vcell counting chamber, as shown in Figure S6.

### 1.5 Detection of cells

After setting the device in the microfluidic flow cell and measuring the CV, cells suspension with a fixed concentration was loaded into the microfluidic chamber (with dimension of 3 mm ×12 mm ×150 μm, corresponding to a volume of 5.4 μL of solution) and incubated (5% CO_2_, 37 °C) for 30 min. After incubation, the cells that were not trapped in the microwells were slightly removed by PBS with a flow of 500 to 1000 μL/mL controlled by a syringe pump. The cells were dyed green or red color with Calcein-AM and monitored by florescence microscopy and CV measurement.

## SI-2 Results and Discussion

**Figure S1.**
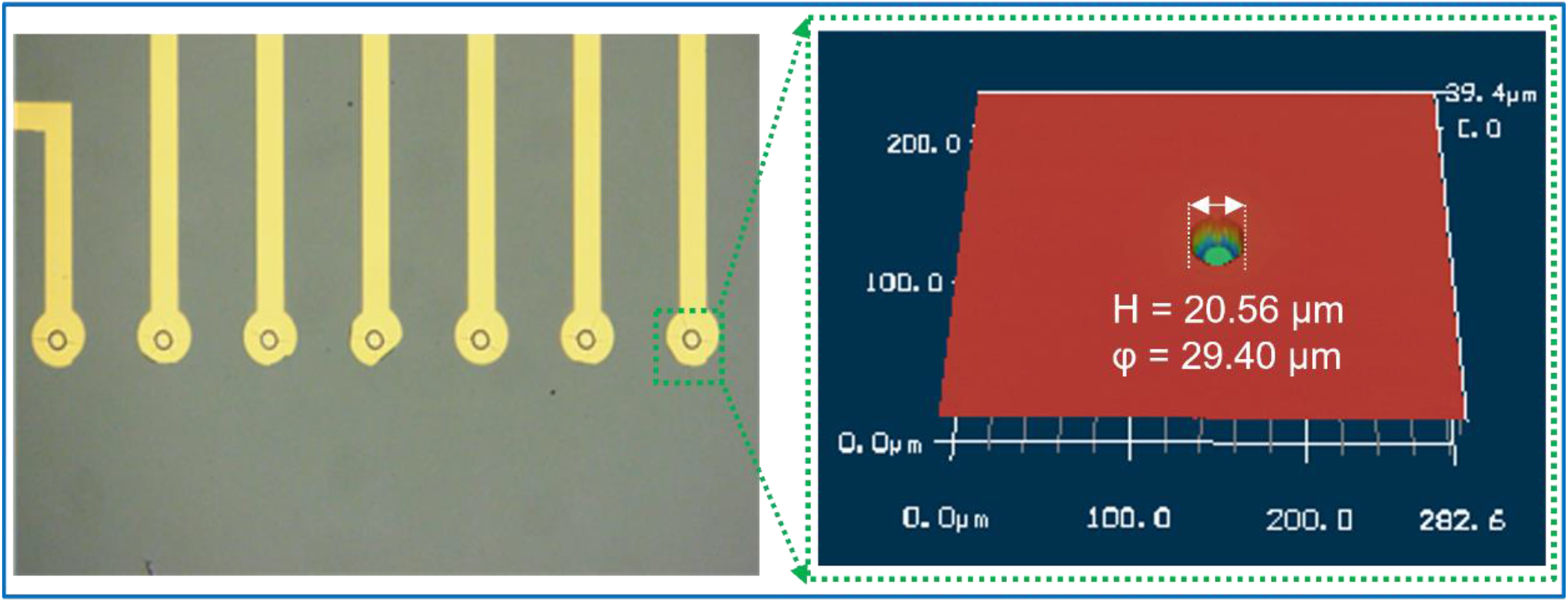
Left: Optical microscope image of part of the electrodes with SU-8 microwells; Right: 3D laser microscope image of single SU-8 Microwell.

**Figure S2.**
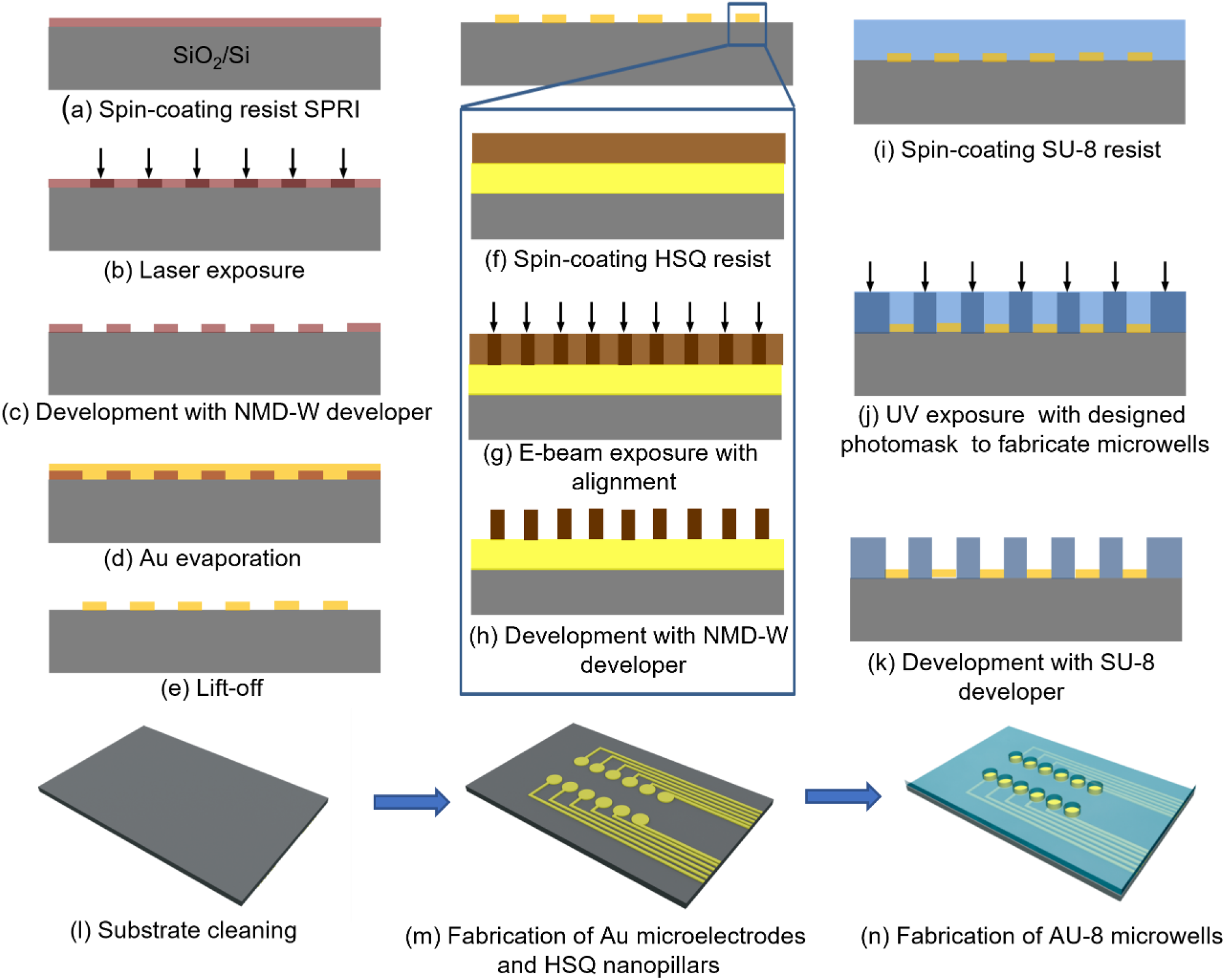
(a) - (e) Au electrodes array fabricated by using Laser lithography. (f) – (h) HSQ nanopillars alignment fabrication process by using e-beam lithography. (i) – (k) SU-8 microwells aligned on the Au electrodes by using UV lithography. (l) – (n) shows the 3D configurations that corresponding to each lithography process.

**Figure S3.**
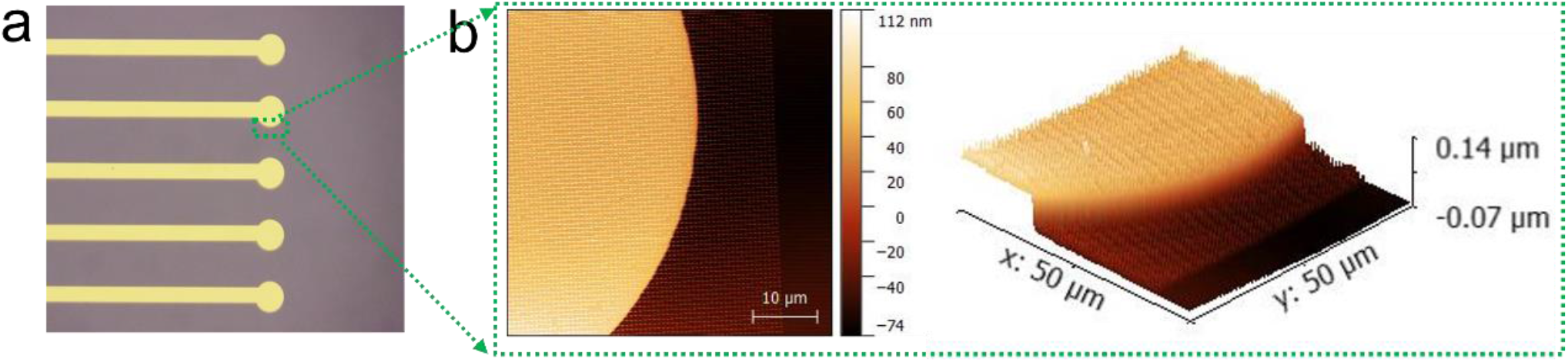
(a) Image of the microelectrode with HSQ nanopillars. (b) 2d and 3d AFM (50 μm x 50 μm) of the HSQ nanopillars aligned on the microelectrode, respectively.

**Figure S4.**
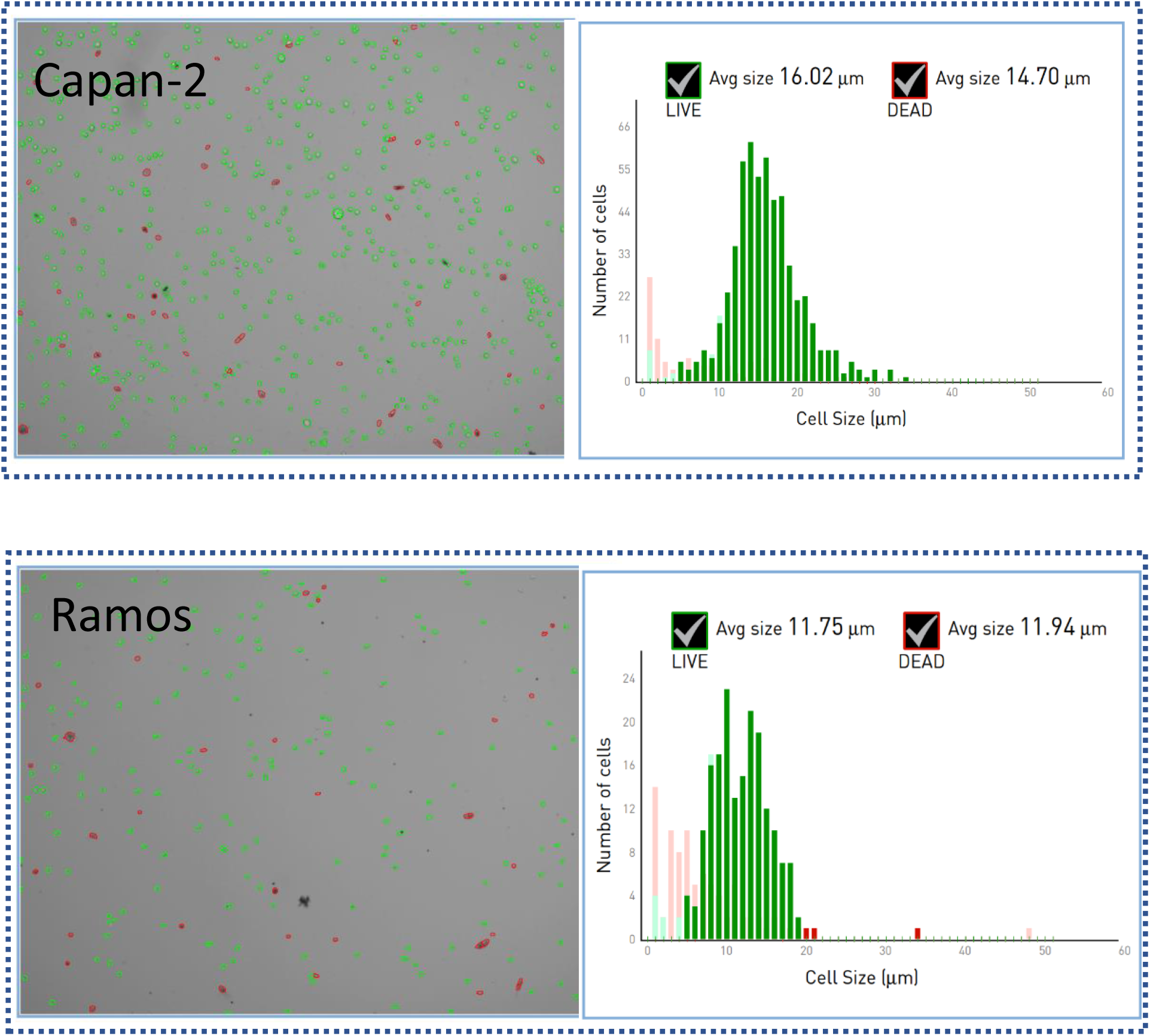
Cell concentration and cell size were determined by using Petroff-Hausser Vcell counting chamber for the two cellular types used in this study: Capan-2 cells and Ramos cells. Green circles correspond to live cells while the red ones correspond to the dead cells.

**Figure S5.**
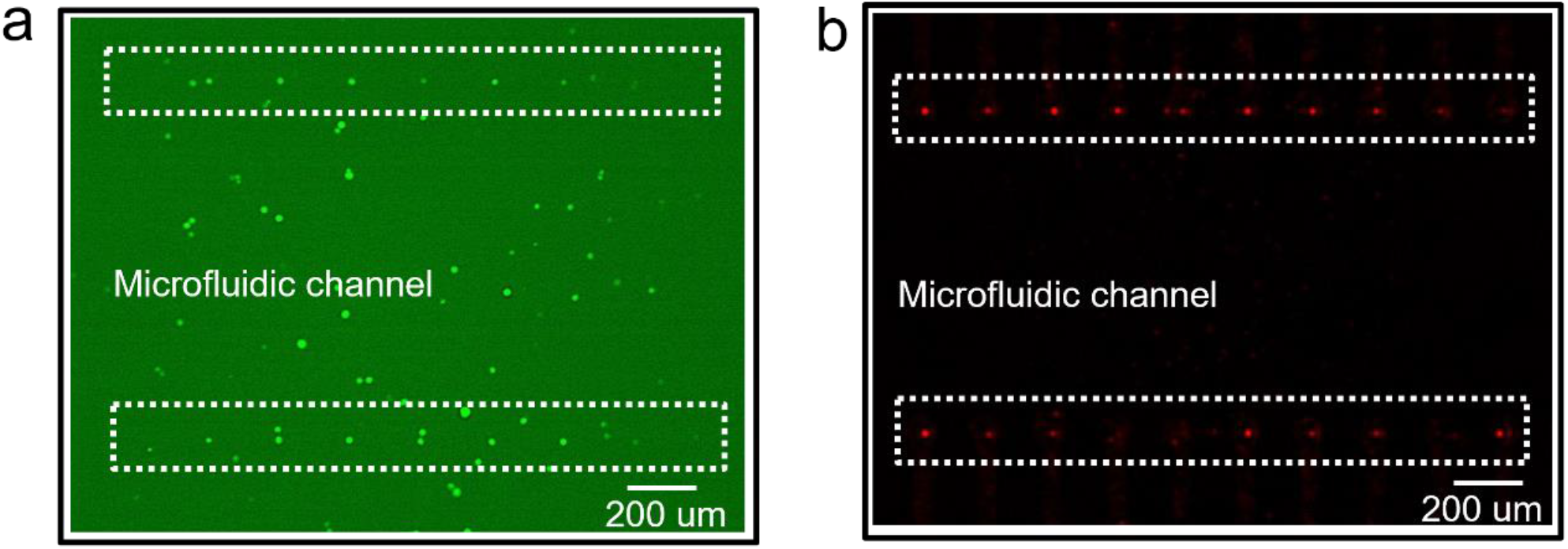
(a) Fluorescence image showing that one chip with 20 devices for single target cell (Capan-2, stained with green colour) trapped on the electrode. (b) Fluorescence image showing that one chip with 20 devices for single non-target cell (Ramos, stained with red colour) trapped on the electrode.

**Figure S6.**
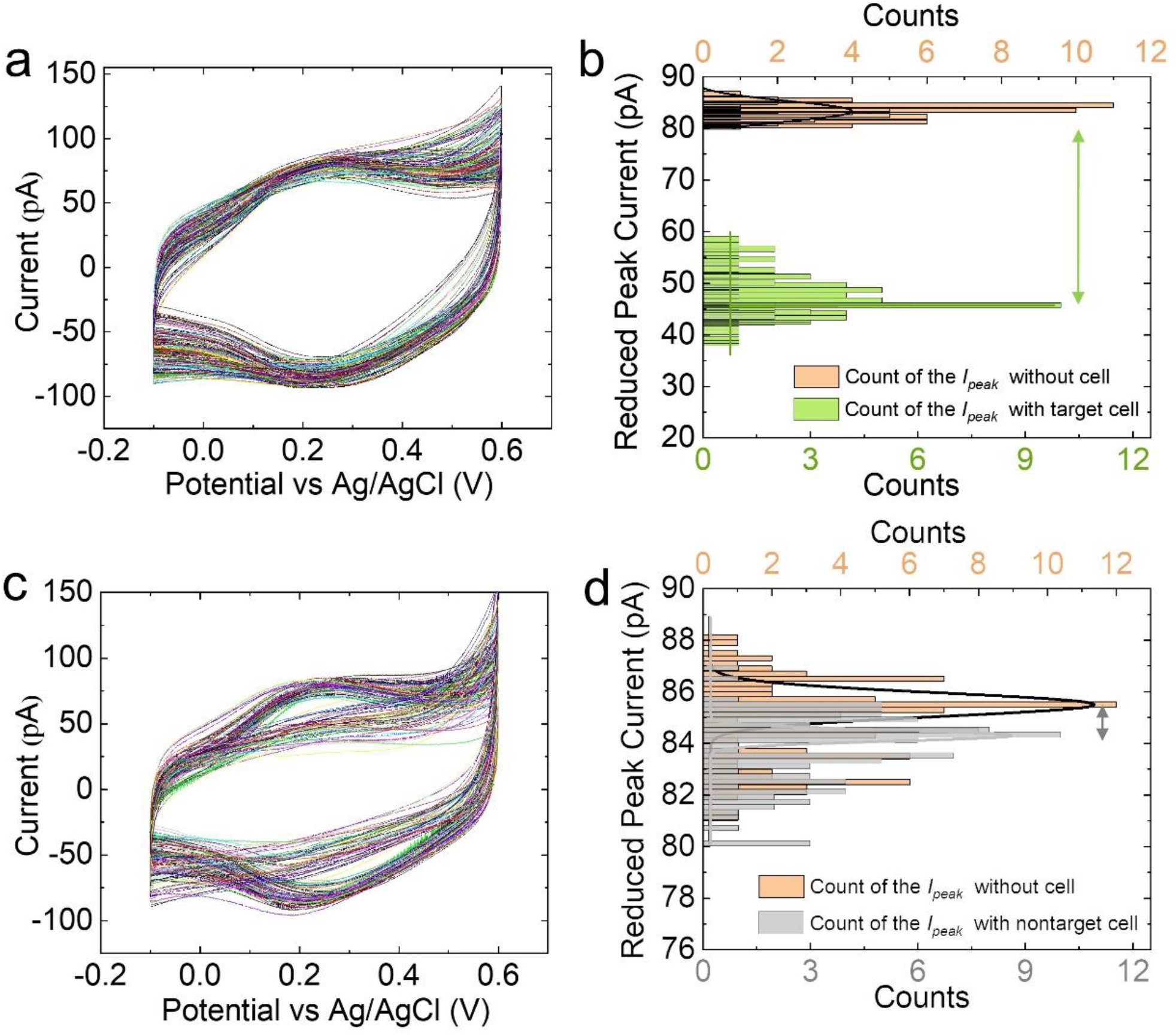
(a) CVs (scan rate *v* = 0.05 V/s) of 100 devices in the presence and absence of single target cell, respectively. (b) Statistics of reduction *I*_*peak*_ of 100 devices for detecting single target cell. (c) CVs (scan rate *v* = 0.05 V/s) of 100 devices in the presence and absence of single non-target cell, respectively. (d) Statistics of reduction *I*_*peak*_ of 100 devices for detecting single non-target cell.

## Notes

### Competing Interest Statement

The authors have declared no competing interest.

### Summary of Updates

This version has been revised based on the review decision letter after submission to a journal

## REFERENCES

(1) Tellez-Gabriel, M.; Ory, B.; Lamoureux, F.; Marie-Francoise Heymann, M. -F.; Heymann, D. Tumour Heterogeneity: The Key Advantages of Single-Cell Analysis. Int. J. Mol. Sci. 2016, 17, 2142.

(2) Cai, L.; Friedman, N.; Xie, X. S. Stochastic protein expression in individual cells at the single molecule level. Nature 2006, 440, 358–362.

(3) Becht, E.; McInnes, L.; Healy, J.; Dutertre, C. A.; W H Kwok, I.; Lai Guan Ng, L. G.; Ginhoux, F.; W Newell, E. Dimensionality reduction for visualizing single-cell data using UMAP. Nat. Biotechnol. 2019, 37 (1) 38–44.

(4) Shapiro, E.; Biezuner, T.; Linnarsson, S. Single-cell sequencing-based technologies will revolutionize whole-organism science. Nat. Rev. Genet 2013, 14, 618–630.

(5) Navin, N.; Hicks, J. Future medical applications of single-cell sequencing in cancer. Genome Med. 2011, 3, 31.

(6) Xu, X.; Hou, Y.; Yin, X.; Bao, L.; Tang, A.; Song, L.; Li, F.; Tsang, S.; Wu, K.; Wu, H.; et al. Single-cell exome sequencing reveals single-nucleotide mutation characteristics of a kidney tumor. Cell 2012, 148, 886–895.

(7) Miwa, H.; Dimatteo, R.; de Rutte, J.; Ghosh, R.; Di carlo, D. Singlecell sorting based on secreted products for functionally defined cell therapies. Microsyst. Nanoeng. 2022, 8, 84.

(8) Kim, S. H., Ito, H., Kozuka, M., Takagi, H., Hirai, M., Fujii, T., Cancer marker-free enrichment and direct mutation detection in rare cancer cells by combining multi-property isolation and microfluidic concentration. Lab Chip 2019, 19 (5), 757–766.

(9) Nagrath, S., Sequist, L. V., Maheswaran, S., Bell, D. W., Irimia, D., Ulkus, L., Smith, M. R., Kwak, E. L., Digumarthy, S., Muzikansky, A., Ryan, P., Balis, U. J., Tompkins, R. G., Haber, D. A., Toner, M., Isolation of rare circulating tumour cells in cancer patients by microchip technology. Nature 2007, 450 (7173), 1235–1239.

(10) Chennit, K., Trasobares, J., Anne, A., Cambril, E., Chovin, A., Clément, N., Demaille, C. Anal. Chem. 2017, 89 (20), 11061–11069.

(11) Mathew, D. G., Beekman, P., Lemay, S. G., Zuilhof, H., Le Gac, S., van der Wiel, W. G. Nano Lett. 2020, 20 (2), 820–828.

(12) Chennit, K., Coffinier, Y., Li, S., Clément, N., Anne, A., Chovin, A. and Demaille, C., High-density single antibody electrochemical nanoarrays. Nano Res. 2022, Nov 18, 1–7.

(13) Zuo, X. L.; Xiao, Y.; and Kevin W. Plaxco, K. W. High Specificity, Electrochemical Sandwich Assays Based on Single Aptamer Sequences and Suitable for the Direct Detection of Small-Molecule Targets in Blood and Other Complex Matrices. J. Am. Chem. Soc. 2009, 131 (20), 6944–6945.

(14) Downs, A. M.; Plaxco. K., W. Real-Time, In Vivo Molecular Monitoring Using Electrochemical Aptamer Based Sensors: Opportunities and Challenges. ACS Sens. 2022, 7 (10), 2823–2832.

(15) Chung, S.; K. Sicklick, J.; Ray, P.; A. Hall, D. Development of a soluble KIT Electrochemical Aptasensor for Cancer Theranostics. ACS Sens. 2021, 6 (5), 1971–1979.

(16) Demaille, C.; Clément, N.; Chovin, A.; Kim, S. H.; Zheng, Z. The Electrochemical Response of Electrode-Attached Redox OligoNucleotides Is Governed by Low Activation Energy Electron Transfer Kinetics. ChemRxiv, December 12, 2022, DOI: 10.26434/chemrxiv-2022-212hs.

(17) Lian, M.; Shi, Y.; Chen, L.; Qin, Y.; Zheng, W.; Zhao, J.; Chen, D. Cell Membrane and V2C MXene-Based Electrochemical Immunosensor with Enhanced Antifouling Capability for Detection of CD44. ACS Sens. 2022, 7 (9), 2701–2709.

(18) Zheng Z.; Kim S., H.; Chovin A.; Demaille, C.; Clément, N. Electrochemical response of surface-attached redox DNA governed by low activation energy electron transfer kinetics. Chem. Sci. 2023, DOI: 10.1039/D3SC00320E.

(19) Grall, S.; Li S.; Jalabert L.; Kim S. H.; Chovin, A.; Demaille C.; Clement, N. Electrochemical Shot-Noise of a Redox Monolayer. arXiv, 2023, DOI: 10.48550/arXiv.2210.12943.

(20) Yang, J. M.; Li, X. D.; Jiang, B. Y.; Yuan, R.; and Yun Xiang. In Situ-Generated Multivalent Aptamer Network for Efficient Capture and Sensitive Electrochemical Detection of Circulating Tumor Cells in Whole Blood. Anal. Chem. 2020, 92 (11), 7893–7899.

(21) Tran, H. L., Dang, V., D. Dega, N., K., Lu, S. -M. Huang, Y. - F. Doong, R., Ultrasensitive detection of breast cancer cells with a lectin-based electrochemical sensor using N-doped graphene quantum dots as the sensing probe. Sens. Actuators B. Chem. 2022, 368, 132233.

(22) Hempel, F.; Law, J. K. Y.; Nguyen, T. C.; Lanche, R.; Susloparova, A.; Vu, X. T.; Ingebrandt, S. PEDOT:PSS organic electrochemical transistors for electrical cell-substrate impedance sensing down to single cells. Biosens. Bioelectron. 2021, 180, 113101.

(23) Bonafè, F.; Decataldo, F.; Zironi, I.; Remmondini, D.; Cramer, T.; Fraboni, B. AC amplification gain in organic electrochemical transistors for impedance-based single cell sensors. Nat. Commun. 2022, 13, 5423.

(24) Shen, H. W.; Deng, W. Q.; He, Y. R.; Li, X. R.; Song, J. L.; Liu, R.; liu, H.; Yang, G. Y.; Li, L. Ultrasensitive aptasensor for isolation and detection of circulating tumor cells based on CeO2@Ir nanorods and DNA walker. Biosens. Bioelectron. 2020, 168, 112516.

(25) Luo, J. J.; Liang, D.; Zhao, D.; Yang, M. H. Photoelectrochemical detection of circulating tumor cells based on aptamer conjugated Cu2O as signal probe, Biosens. Bioelectron. 2020, 151, 111976.

(26) Li, S.; Coffinier, Y.; Lagadec, C.; Cleri, F.; Nishiguchi, K.; Fujiwara, A.; Fujii, T.; Kim, S. H.; Clement, N. Redox-labelled electrochemical aptasensors with nanosupported cancer cells. Biosens. Bioelectron. 2022, 216, 114643.

(27) Song, Y.; Zhu, Z.; An, Y.; Zhang, W.; Zhang, H.; Liu, D. Yu, C.; Duan, W.; Yang, C. J. Selection of DNA Aptamers against Epithelial Cell Adhesion Molecule for Cancer Cell Imaging and Circulating Tumor Cell Capture. Anal. Chem. 2013, 85 (8), 4141–4149.

(28) Clement, N.; Patriarche, G.; Smaali, K.; Vaurette, F.; Nishiguchi, K.; Troadec, D.; Fujiwara, A.; Vuillaume, D. Large array of sub-10-nm single-grain Au nanodots for use in nanotechnology. Small 2011, 7 (18), 2607–2613.

(29) Trasobares, J.; Rech, J.; Jonckheere, T.; Martin, T.; Aleveque, O.; Levillain, E.; Diez-Cabanes, V.; Olivier, Y.; Cornil, J.; Nys, J. P.; Sivakumarasamy, R.; Smaali, K.; Leclere, P.; Fujiwara, A.; Theron, D.; Vuillaume, D.; Clement, N. A 17 GHz molecular rectifier. Nat. Commun. 2016, 7, 12850.

(30) Anne, A.; Demaille, C. Dynamics of Electron Transport by Elastic Bending of Short DNA Duplexes. Experimental Study and Quantitative Modeling of the Cyclic Voltammetric Behavior of 3’-Ferrocenyl DNA End-Grafted on Gold. J. Am. Chem. Soc. 2006, 128, 542–557.

(31) Steentjes, T.; Jonkheijm, P.; Huskens, J. Electron Transfer Processes in Ferrocene-Modified Poly(ethylene glycol) Monolayers on Electrodes. Langmuir 2017, 33 (43), 11878–11883.

(32) Dauphin-Ducharme, P.; Arroyo-Currás, N.; Adhikari, R.; Somerson, J.; Ortega, G.; Makarov, D. E.; Plaxco, K. W. Chain Dynamics Limit Electron Transfer from Electrode-Bound, Single Stranded Oligonucleotides. J. Phys. Chem. C 2018, 122 (37), 21441–21448.

(33) Xie, X.; Li, F.; Zhang, H.; Lu, Y.; Lian, S.; Lin, H.; Gao, Y.; Jia, L. EpCAM aptamer-functionalized mesoporous silica nanoparticles for efficient colon cancer cell-targeted drug delivery. Eur. J. Pharm. Sci. 2016, 83, 28–35.

(34) Mayado, A.; Orfao, A.; Mentink, A.; Gutierrez, M. L.; Muñoz-Bellvis, L. Terstappen, L. W. M. M. Detection of circulating tumor cells in blood of pancreatic ductal adenocarcinoma patients. Cancer Drug Resist. 2020, 3, 83–97.

(35) Grall, S., Alic, I., Pavoni, E., Awadein, M., Fujii, T., Mullegger, S., Farina, M., Clément, N., Gramse, G. Attoampere nanoelectrochemistry. Small 2021, 17 (29), 2101253.

(36) Kutovyi, Y., Hlukhova, H., Boichuk, N., Menger, M., Offenhäusser, A., Vitusevich, S., Amyloid-beta peptide detection via aptamer-functionalized nanowire sensors exploiting single-trap phenomena. Biosens. Bioelectron. 2020, 154, 11205.

## Supplementary References

(1) Zhou, J.; Zhang, X.; Sun, J.; Dang, Z.; Li, J.; Li, X.; Chen, T. The effects of surface topography of nanostructure arrays on cell adhesion. Phys. Chem. Chem. Phys. 2018, 20 (35), 22946–22951.

(2) Li, S.; Coffinier, Y.; Lagadec, C.; Cleri, F.; Nishiguchi, K.; Fujiwara, A.; Fujii, T.; Kim, S. H.; Clement, N. Redox-labelled electrochemical aptasensors with nanosupported cancer cells. Biosens. Bioelectron. 2022, 216, 114643.

